# Single-cell molecular profiling provides a high-resolution map of basophil and mast cell differentiation

**DOI:** 10.1101/848358

**Authors:** Fiona K. Hamey, Winnie W.Y. Lau, Iwo Kucinski, Xiaonan Wang, Evangelia Diamanti, Nicola K. Wilson, Berthold Göttgens, Joakim S. Dahlin

## Abstract

Differentiation of hematopoietic stem and progenitor cells ensure a continuous supply of mature blood cells. Recent models of differentiation are represented as a landscape, in which individual progenitors traverse a continuum of multipotent cell states before reaching an entry point that marks lineage commitment. Basophils and mast cells have received little attention in these models and their differentiation trajectories are yet to be explored. Here, we have performed multicolor flow cytometry and high-coverage single-cell RNA sequencing analyses to chart the differentiation of hematopoietic progenitors into basophils and mast cells in mouse. Analysis of flow cytometry data reconstructed a detailed map of the differentiation, including a bifurcation of progenitors into two specific trajectories. Molecular profiling and pseudotime ordering of the single cells revealed gene expression changes during differentiation, with temporally separated regulation of mast cell protease genes. We validate that basophil and mast cell signature genes increased along the trajectories into their respective lineage, and we demonstrate how genes critical for each respective lineage are upregulated during the formation of the mature cells. Cell fate assays showed that multicolor flow cytometry and transcriptional profiling successfully predict the bipotent phenotype of a previously uncharacterized population of basophil-mast cell progenitor-like cells in mouse peritoneum. Taken together, we provide a detailed roadmap of basophil and mast cell development through a combination of molecular and functional profiling.

## Introduction

Hematopoietic stem and progenitor cells (HSPCs) constitutively generate blood cells, including erythrocytes, platelets, granulocytes, macrophages, and lymphocytes. The hierarchical model of hematopoiesis, with distinct megakaryocyte-erythroid, granulocyte-monocyte, and lymphoid branches, was the dominating representation of the differentiation process before the introduction of single-cell RNA sequencing.^1^ Recent groundbreaking studies that couple single-cell transcriptomics and cell fate assays reveal that blood cell differentiation more likely represents a landscape of cell states with continuous progression from progenitors into the mature cell lineages.^2–6^

Mast cells and basophils constitute two cell types of the hematopoietic system, whose differentiation trajectories are yet to be deciphered. Mast cells are sentinel cells that are strategically positioned throughout the body and allow rapid triggering of the immune system upon infection.^7^ Along with basophils, their activation results in prompt release of proteases and histamine from the cytoplasmic granules as well as synthesis of cytokines and chemokines. These mediators in turn cause inflammation, vasodilation, and leukocyte recruitment to the site of triggering.^7^ A similar cascade is initiated following IgE-allergen-mediated cell activation that causes allergic symptoms in patients.

Bone marrow HSPCs give rise to basophil and mast cells,^8^ and single-cell transcriptomics of Lin^−^ c-Kit^+^ mouse bone marrow progenitors recently uncovered the gene expression changes during the transition from hematopoietic stem cells to common bipotent basophil-mast cell progenitor (BMCP).^3^ However, the further progression of BMCPs to basophils and mast cells is yet to be delineated.

Here, we combine multicolor FACS index sorting with high-coverage single-cell RNA sequencing to investigate the basophil-mast cell bifurcation and the differentiation into each respective lineage. We demonstrate that molecular profiling and pseudotime ordering of single cells highlights genes that are critical for cell differentiation and maturation. The analysis is accompanied with the generation of a user-friendly web resource that allows gene expression to be explored across the single-cell landscape. Finally, we use cell-fate assays to show that single-cell transcriptomics and protein epitope data analysis successfully predict the fate potential of the previously uncharacterized BMCP-like cell population in the peritoneal cavity. Taken together, the current resource provides a detailed roadmap of two rare and developmentally related hematopoietic cells, whose activation contributes to a broad range of human diseases.

## Methods

### Cell isolation and flow cytometry

Experiments involving mice were performed according to the United Kingdom Home Office regulations. PBS with 2 % fetal calf serum (Sigma-Aldrich, St Louis, MO) and 1 mM EDTA was injected into the peritoneal cavity of euthanized C57BL/6 mice. The fluid was aspirated following vigorous massage, and the cells were prepared for fluorescence-activated cell sorting (FACS). Peritoneal lavage samples with excessive blood contamination were discarded before data acquisition. Bone marrow cells were extracted by flushing or crushing the femurs, tibias, and/or ilia. Red blood cells were lysed and the remaining cells were prepared for FACS. The cells were sorted with a BD Influx cell sorter (BD Biosciences, San Jose, CA). Cell doublets were excluded with the width parameters. BMCP-like cells and mast cells were sorted two consecutive times for cell culture experiments. The cells were sorted into Terasaki plates (Greiner Bio-One, Kremsmünter, Austria) or 96-well plate wells. Visual inspection determined colony sizes following culture, and the size was set to 1 if no live cells were observed in a particular well. Flow cytometry was typically performed on colonies constituting at least 20 cells as described previously.^3^ Cultured cells were analyzed with the BD Fortessa flow cytometers (BD Biosciences).

### Antibodies and cell staining

Primary cells were incubated with the antibodies mouse hematopoietic progenitor cell isolation cocktail, integrin β7 (clone FIB504), CD34 (RAM34), Sca-1 (D7), CD16/32 (93), c-Kit (2B8), FcεRI (MAR-1), IL-33Rα/ST2 (DIH9), and/or CD49b (DX5). Cultured cells were stained with c-Kit, FcεRI, CD49b, with or without TER119 (TER119). Fc-block (clone 93) was used where appropriate. The antibodies were from STEMCELL Technologies (Vancouver, Canada), BD Biosciences, Biolegend (San Diego, CA), and Thermo Fisher Scientific (Waltham, MA). DAPI (BD Biosciences) or 7-AAD (Thermo Fisher Scientific) were used to exclude dead cells.

### Cell culture

The cells were cultured for 6-7 days in IMDM (Sigma-Aldrich) with 20 % heat-inactivated fetal calf serum (Sigma-Aldrich), 100 U/ml penicillin (Sigma-Aldrich), 0.1 mg/ml streptomycin (Sigma-Aldrich), 50-200 μM β-mercaptoethanol (Thermo Fisher Scientific). The medium was supplemented with 20 ng/ml IL-3 and 100 ng/ml stem cell factor, or 80 ng/ml stem cell factor, 20 ng/ml IL-3, 50 ng/ml IL-9, and 2 U/ml erythropoietin. All cytokines were recombinant mouse cytokines (Peprotech, Rocky Hill, NJ) except the erythropoietin (Eprex; Janssen-Cilag, High Wycombe, UK), which was human.

### Flow cytometry analysis

FlowJo v10 (Treestar, Ashland, OR) produced the flow cytometry plots. Diffusion map and principal component analysis plots of flow cytometry data were generated using the R programming environment. Briefly, the flow cytometry events were down-sampled according to the population with the least number of events. Duplicate entries were removed, and the parameters representing fluorescent markers log-transformed. Variables were z-scored and diffusion map plots generated using the *destiny* and *ggplot2* packages. Principal component analysis (PCA) was calculated using the prcomp function. Data projection was performed using the predict function.

### Single-cell RNA sequencing and analysis

Single-cells were FACS index sorted into lysis buffer, and single-cell RNA sequencing was performed based on the Smart-Seq2 protocol.^9^ Protocols and single-cell RNA sequencing data generated for this article have been deposited in the Gene Expression Omnibus database (accession numbers GSE128003 and GSE128074). Single-cell RNA sequencing data of bone marrow BMCPs, analyzed in Dahlin et al (2018),^3^ are available through GSE106973.

Sequencing data were aligned using GSNAP^10^ to Ensembl genome build 81^11^ and gene counts were obtained using HT-Seq^12^. Quality control filtering and normalization was performed in the R programming environment. Quality control was performed to exclude cells with fewer than 500,000 reads mapping to nuclear genes or with over 25% of mapped reads mapping to ERCC spike-ins. For the peritoneal cells, PCA of the quality control-filtered samples showed that two cells separated from the rest of the sample in principal component (PC) 1 (Figure S2A). Genes with high PC1 loadings were highly significantly enriched for B cell related genes, so these two outlier cells were suspected to be contaminating B cells and excluded from further analysis. Cells were then normalized using Scran^13^ and highly variable genes were identified using the ERCC spike-ins to estimate technical variance^14^. This identified 3330 highly variable genes for the basophil dataset and 1832 highly variable genes for the mast cell dataset.

Downstream analysis was performed using the scanpy v1.4 python module^15^. Inbuilt scanpy functions were used for PCA and diffusion map dimensionality reduction. Differential expression between cell types was performed using the *rank_genes_groups* function with the t-test_overestim_var option for testing. P-values were adjusted using the benjamini-hochberg method for correcting for multiple testing, and genes with adjusted p-value < 0.01 were considered significant. Gene list enrichment analysis was performed using the *enrichr* function from the *gseapy* python module^16, 17^. Cell cycle scoring was performed on scaled data using the scanpy *score_genes_cell_cycle* function with S phase and G2/M phase gene lists downloaded from Macosko et al^18^. Bone marrow BMCP cells from Dahlin et al (2018)^3^ were projected into the PCA space of the peritoneal cells and the k=10 closest peritoneal neighbors of each bone marrow cell were identified in these co-ordinates. Figure 3B was then generated by scoring how frequently each peritoneal cell was the nearest neighbor of a bone marrow BMCP. Peritoneal mast cells were ordered in pseudotime using the diffusion pseudotime (DPT) scanpy implementation^19^. Due to cell cycle effects confounding the diffusion map and DPT analysis, basophil progenitor cells were ordered in pseudotime by ordering cells along PC1. Genes with dynamic expression in pseudotime were identified following the method of Tusi et al.^2^ Briefly, gene expression was first smoothed along pseudotime using a sliding window of size 20. For each ordering, the windows with minimum and maximum gene expression were identified, and a t-test performed between the values in each of these windows, giving a p-value for each gene. To generate a background distribution, this analysis was repeated for a random shuffling of cells along pseudotime. The adjusted p-value for each gene was then calculated as the fraction of shuffled p-values across all genes that were less than the p-value of the gene in question for non-permuted data. Genes with adjusted p-value < 0.01 were then treated as dynamic across pseudotime and plotted in the heatmaps. Gene expression in the heatmap was smoothed using a sliding window of size 20 and z-score transformed for each gene. To identify groups of genes with different pseudotime dynamics, genes were clustered using Louvain clustering^20^ with the scanpy implantation, with the nearest neighbor matrix calculated on the full pseudotime expression matrix. The resolution was chosen to obtain 2 clusters for each dataset: downregulated and upregulated genes. Mast cell and basophil signature gene sets were obtained from Dwyer et al,^21^ who used microarray analysis on bulk samples to characterize gene expression specific to these mature cell types. Statistical overlap between gene lists was calculated using a hypergeometric test. Panther v14.1 was used to identify the dynamic mast cell genes annotated as proteases (Panther category PC00190)^22^. To plot gene expression trends along pseudotime the genSmooth Curves function from the monocle R package^23^ was used to fit smooth spline curves for the expression of each gene against pseudotime. When all genes were plotted together expression values of each gene were scaled by dividing values by the maximum of that gene along pseudotime to account for the very different dynamic ranges across genes. Interactive websites for plotting gene expression and flow cytometry data are hosted at http://128.232.224.252/bas/ and http://128.232.224.252/per/ for the basophil and mast cell dataset, respectively.

## Results

### Multicolor flow cytometry analysis reveals the basophil and mast cell differentiation trajectories

Basophil and mast cell differentiation are closely linked, and the cells share a common bipotent progenitor (Figure 1A). Here, we used multicolor flow cytometry to map these branching trajectories at the single-cell level. Flow cytometry analysis of mouse bone marrow cells captured BMCPs and cells of the basophil differentiation trajectory (Figure 1Bi,ii).^3, 24^ As the late mast cell differentiation takes place at peripheral sites, parallel analysis of peritoneal cells identified BMCP-like cells and mast cells (Figure 1Biii). Dimensionality reduction with a diffusion map algorithm enabled 2-dimensional visualization of the flow cytometry single-cell datasets, which covered 5 cell populations recorded with 9 fluorescent and 2 light scatter parameters. The diffusion map visualization revealed a bifurcation at the BMCP stage, establishing the putative entry points to the basophil and mast cell trajectories (Figure 1C). The diffusion map embedding further visualized the progression from BMCP, through basophil progenitors, to basophils. The mast cell trajectory exhibited a similar pattern, with differentiation of BMCP-like cells to mature mast cells.

**Figure 1.**
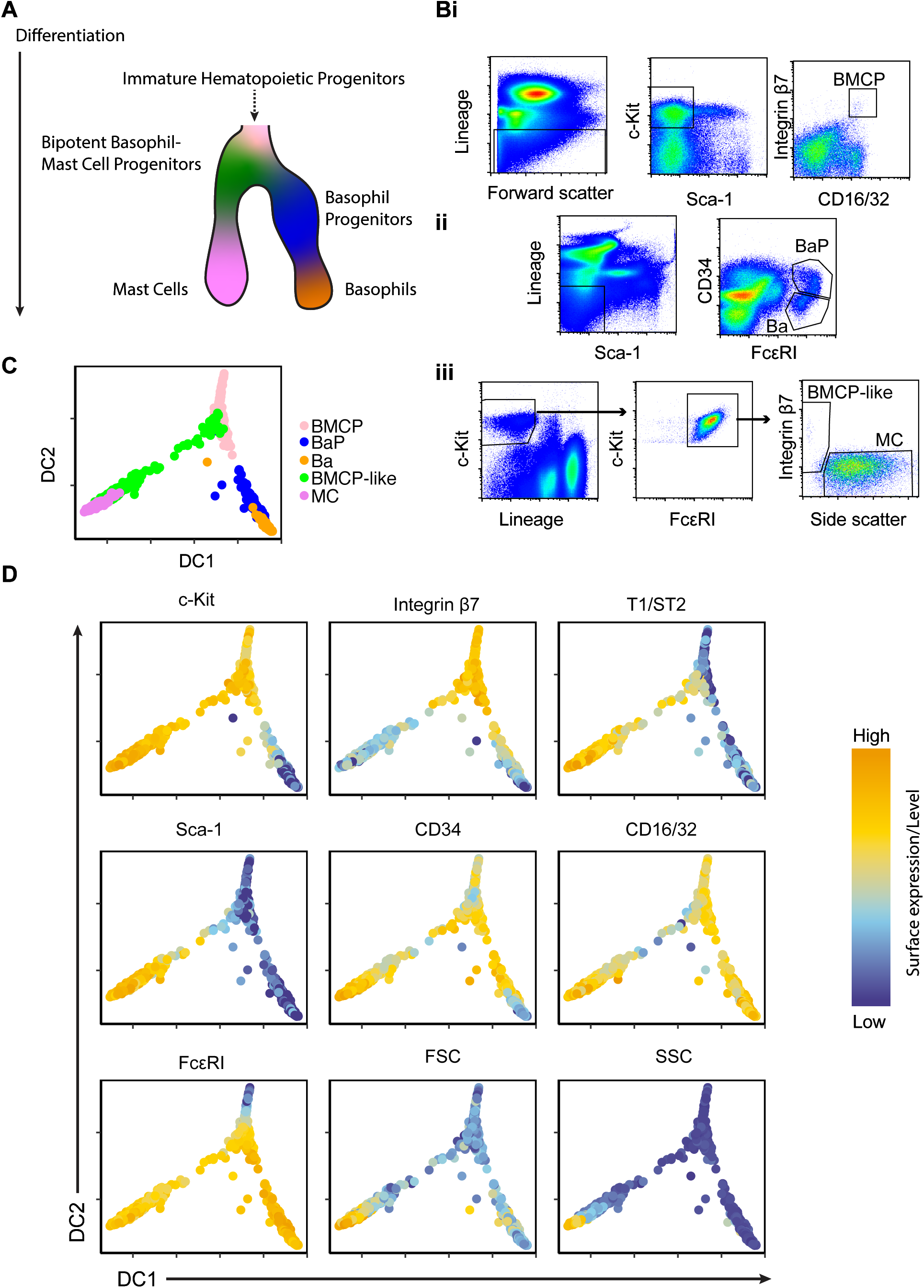
Flow cytometry analysis reveals differentiation trajectories from bipotent basophil-mast cell progenitors to basophils and mast cells. (A) Illustration outlining the basophil and mast cell differentiation trajectories. (B) Flow cytometry-based gating strategies of (Bi) bipotent basophil-mast cell progenitors (BMCPs) from bone marrow, (Bii) basophil progenitors (BaP) and basophils (Ba) from bone marrow, and (Biii) BMCP-like cells and mast cells from peritoneal cavity. (C) Diffusion map visualization of the flow cytometry data colored by cell type. (D) Diffusion map visualization of the flow cytometry data colored by protein expression or light scatter parameters. The surface expression parameters and light scatter parameters are visualized on log-transformed and linear scales, respectively. Expression of lineage markers and viability staining are not shown. The data are representative of 4 independent experiments.

Plotting individual surface markers in the diffusion map allowed us to investigate how the proteins are expressed during differentiation. For example, loss of CD34 in combination with downregulation of c-Kit marked the progression from BMCPs to basophils (Figure 1D), and loss of integrin β7 in c-Kit^+^ cells was associated with differentiation along the trajectory from BMCPs to mast cells (Figure 1D). Taken together, the flow cytometry dataset provides a template of basophil-mast cell differentiation at single-cell level and highlights the bifurcation towards the two lineages.

### Single-cell profiling captures progression of basophil differentiation in the bone marrow

Analysis by flow cytometry suggested that the flow cytometry gating strategies we used could be capturing a continuum of differentiation towards basophils and mast cells. To first identify changes in gene expression programs during basophil differentiation, we performed single-cell RNA-sequencing of basophil progenitor (BaP) cells and basophil (Ba) cells from mouse bone marrow. Both principal component analysis (PCA) and diffusion maps showed separation between the majority of cells from the two sorting gates (Figure 2A, Figure S1A). To investigate which genes were driving this separation, we performed differential gene expression analysis, identifying 212 genes upregulated and 833 genes downregulated in Ba cells compared to BaPs (Table S1). Enrichment analysis of these gene lists revealed that upregulated genes were enriched for granulocyte immune response terms (Figure S1B). Downregulated genes were enriched for cell cycle related terms (Figure 2B), suggesting a difference in cell cycle behavior throughout the differentiation process. This observation is in line with other hematopoietic differentiation pathways, where progenitors commonly loose proliferative capacity as they mature into the fully differentiated cell types.

**Figure 2.**
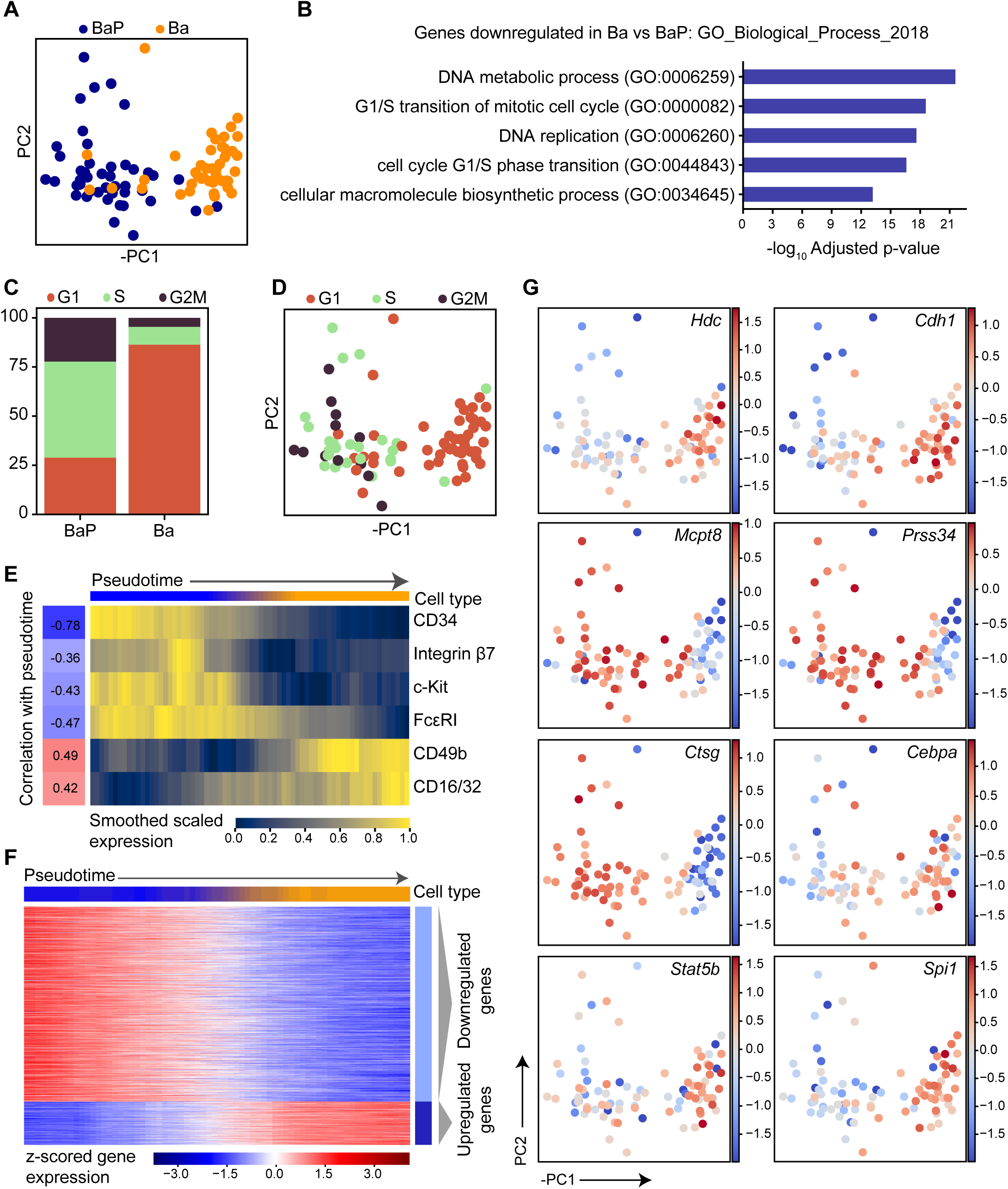
Bone marrow basophil progenitors downregulate cell cycle genes during differentiation. (A) PCA of scRNA-seq profiles colored by cell surface marker phenotype. PC, principal component. (B) Top 5 GO Biological Process terms associated with the genes significantly upregulated in BaP cells compared to Ba cells, ranked by adjusted p-value. Benjamini-Hochberg correction for multiple hypotheses testing. Genes upregulated in Ba compared to BaP are presented in Figure S1B. (C) Proportion of scRNA-seq profiles from each phenotype computationally assigned to G1, S or G2M cell cycle states based on gene expression. (D) PCA colored by cell cycle state. (E) Levels of cell surface markers for cells ordered by PC1 pseudotime. Index data values were log-transformed, smoothed along pseudotime by using a sliding window of size 20 and scaled between 0 and 1 for each marker. Correlation values indicate the pearson correlation coefficient between pseudotime and the unsmoothed expression values for each surface marker. Colorbar at the top indicates the phenotypic cell type proportions within each window. Blue corresponds to entirely BaPs and orange to Ba cells. (F) Heatmap displaying the expression of genes dynamically expressed along the PC1 pseudotime ordering. The top colorbar indicates the cell type proportion in each window. Expression is smoothed along a sliding window and z-scored for each gene, and genes were clustered using Louvain clustering into groups showing different dynamics. (G) PCA colored by z-scored expression of specific genes. The data represents cells pooled from 4 individual mice.

To further explore this, we then performed analysis to computationally assign cell cycle state to the single-cell profiles.^18^ Consistent with the gene list enrichment analysis, the majority of cells in the BaP gate were assigned to S and G2M states (69%), whereas 87% of cells in the Ba gate were assigned to G1 state (Figure 2C, D). The effect of cell cycle status was clear in the diffusion map dimensionality reduction (Figure S1C), confounding attempts to order cells using pseudotime algorithms such as diffusion pseudotime (DPT). Instead, downregulation of progenitor marker genes such as *Cd34* and *Kit* indicated that ordering cells along the first principal component (PC) could be used to arrange cells in pseudotime (Figure S1D). Visualization of cell surface markers measured by index sorting also showed clear dynamics of the different surface markers along PC1 (Figure 2E). As expected, CD34 and c-Kit protein expression showed a negative correlation with pseudotime (compare Figure 1D and 2E), which indicates their downregulation during basophil differentiation. In addition, the basophil marker CD49b (DX5) showed a positive correlation with pseudotime ordering (Figure 2E).

Using the PC1 pseudotime ordering, we then identified genes that dynamically changed during differentiation (Figure 2F). Clustering sorted these dynamic genes into two groups: one increasing and one decreasing with differentiation (Table S2). Basophil differentiation was associated with upregulation of *Hdc*, which is associated with histamine synthesis, and increased expression of the basophil gene E-cadherin (*Cdh1*). We further observed downregulation of the proteases *Mcpt8*, *Prss34* and *Ctsg* and upregulation of the transcription factors *Cebpa*, *Stat5b*, and *Spi1* (Figure 2G). To validate the full lists of dynamically regulated genes, we compared these to mast cell and basophil signature genes identified using bulk microarray analysis.^21^ Genes upregulated during basophil differentiation exhibited a significant overlap with the previously described basophil signature genes (p = 4.0 × 10^-29^, hypergeometric test, Figure S1Ei), whereas genes that were downregulated during differentiation had significant overlap with the previously described mast cell signature genes (p = 1.3 × 10^-4^, hypergeometric test, Figure S1Eii).

Together, this analysis offers a description of the dynamics of gene expression during basophil differentiation and highlights changes in cell cycle activity as one of the major occurrences during this maturation process.

### Single-cell gene expression analysis suggests a continuum of mast cell differentiation in the peritoneal cavity

After exploring the basophil progenitors, we next decided to focus on mast cell differentiation in the peritoneal cavity. The flow cytometry data suggested the existence of both BMCP-like peritoneal cells and peritoneal mast cells (Figure 1), so we performed single-cell RNA-sequencing on these populations to characterize them based on gene expression. A subset of the BMCP-like cells clustered separately from the mast cells in the diffusion map plot, demonstrating a difference between the transcriptome of these cells and the peritoneal mast cells (Figure 3A). In previous work we characterized bone marrow BMCPs at the single-cell gene expression level.^3^ To examine the similarity of these bone marrow progenitors to the peritoneal mast cell differentiation, single-cell bone marrow BMCP profiles from Dahlin et al^3^ were projected onto the peritoneal dataset (Figure 3B). This demonstrated that the BMCP-like peritoneal cells furthest from the peritoneal MCs were most similar to the bone marrow BMCPs, supporting that these were the most immature cells in the dataset.

**Figure 3.**
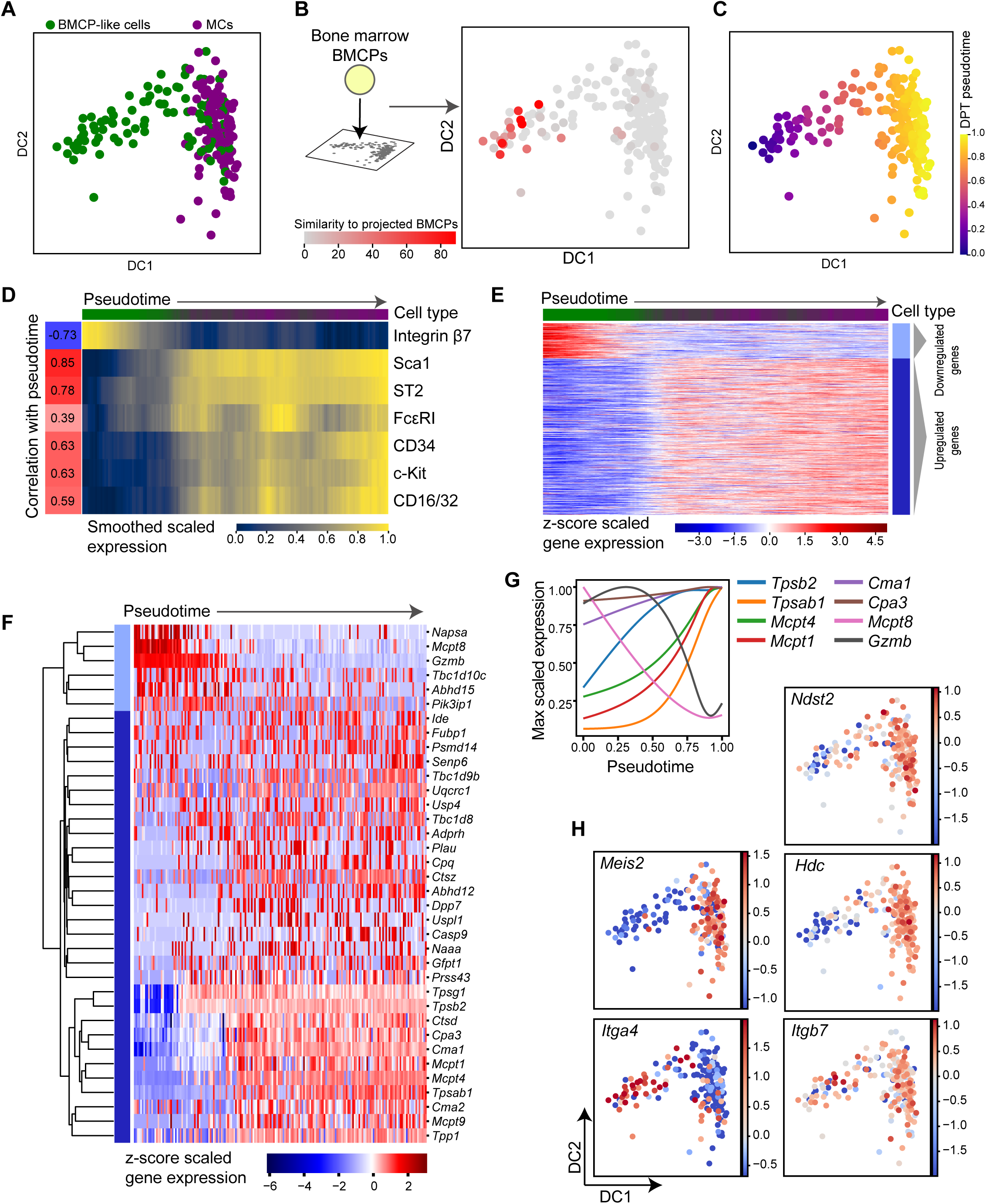
Transcriptional profiling of peritoneal mast cell progenitors captures a differentiation continuum. (A) Diffusion map dimensionality reduction of scRNA-seq profiles colored by cell phenotype. DC, diffusion component. (B) Projection of bone marrow BMCP progenitor scRNA-seq profiles from Dahlin et al (2018)^3^ to their most similar expression profiles from the peritoneal dataset. Each peritoneal cell is colored by its similarity to the projected bone marrow cells, see methods for details. (C) Diffusion map colored by pseudotime ordering of cells. DPT, diffusion pseudotime. (D) Levels of cell surface markers for pseudotime ordered cells. Index data values were log-transformed, smoothed along pseudotime by using a sliding window of size 20 and scaled between 0 and 1 for each marker. Correlation values indicate the pearson correlation coefficient between pseudotime and the unsmoothed expression values for each surface marker. Colorbar at the top indicates the phenotypic cell type proportions within each window. Green corresponds to entirely BMCP-like cells and purple to MCs. (E) Heatmap displaying the expression of genes dynamically expressed along the pseudotime ordering. The top colorbar indicates the proportion of cell type in each window. Expression is smoothed along a sliding window and z-scored for each gene, and genes were clustered using Louvain clustering into groups showing different dynamics. (F) Heatmap of dynamically regulated proteases showing z-scored gene expression along pseudotime. Genes were ordered using the hierarchical clustering indicated by the dendrogram. Colorbar indicates the Louvain cluster from (E) for each gene. (G) Expression trends of specific genes along pseudotime. Genes are scaled by their maximum expression value rather than z-scoring as in the heatmap. (H) Diffusion map colored by z-score scaled expression of specific genes. The data represents cells pooled from 3 individual mice.

To understand expression changes during mast cell maturation, we then performed pseudotime ordering of the peritoneal cells using DPT (Figure 3C). As expected, interrogation of cell surface markers along the pseudotime ordering showed a strong downregulation of integrin β7 and strong upregulation of markers such as Sca1 and ST2 (compare Figure 1D and 3D). Genes exhibiting dynamic expression patterns were identified and clustered as for the basophil trajectory (Table S3, Figure 3E). Annotation from the Panther database^22^ was used to interrogate the two gene clusters for overlap with specific annotated gene sets such as proteases. Protease genes downregulated during mast cell differentiation included *Mcpt8* and *Gzmb*, whereas *Cpa3*, *Cma1*, *Mcpt1*, *Mcpt4*, *Tpsb2,* and *Tpsab1* increased with differentiation (Figure 3F). To investigate the temporal induction and loss of protease genes, we changed visualization method and scaled the gene expression according to the cell with maximum expression (instead of z-scoring genes across the dataset). Plotting the maximum value-scaled gene expression revealed the gene dynamics across the pseudotime trajectory. Early onset proteases included *Cpa3*, followed by *Tpsb2*, and finally *Tpsab1*, indicating that the protease induction occurs in stages (Figure 3G, raw values for individual genes shown in Figure S2C).

To validate the full lists of dynamically regulated genes in the peritoneal mast cell dataset, we compared these to mast cell and basophil signature identified in Dwyer et al.^21^ The upregulated genes significantly overlapped with the mast cell signature genes (p = 3.7 × 10^-65^, hypergeometric test, Figure S2Di). Upregulated genes included *Ndst2*, *Meis2* and *Hdc* (Figure 3H). Some genes showed expression enrichment mainly in the mast cells (*Meis2*), whereas others were expressed more evenly across the trajectory except for lower expression at the beginning of pseudotime (*Ndst2*). There was also a small overlap between the downregulated genes and basophil signature genes (p = 2.5 × 10^-5^, hypergeometric test, Figure S2Dii). To investigate the link between gene and protein expression we also interrogated the expression of *Itga4* and *Itgb7*, which encode subunits of Integrin β7. *Itga4* was significantly downregulated with a similar expression pattern to integrin β7 in the flow cytometry data whereas *Itgb7* was not significantly dynamically changing in pseudotime (Figure 3D, H).

### BMCP-like cells in the peritoneal cavity exhibit basophil and mast cell-forming potential

The flow cytometry-based and transcriptional analyses revealed an immature cell population with BMCP-like characteristics in the peritoneal cavity. However, a population of bipotent peritoneal BMCPs has not previously been described at this site. We therefore explored whether the protein and transcriptional analyses successfully predicted the developmental state of the peritoneal BMCP-like cells and mast cells. FACS-isolated BMCP-like cells and mast cells were cytochemically stained with May-Grünwald Giemsa. Primary BMCP-like cells displayed little cytoplasm that contained no or few granules, consistent with the morphology of blasts (Figure 4A). In contrast, primary mast cells were filled with numerous metachromatic granules, in agreement with a mature morphology (Figure 4A).

**Figure 4.**
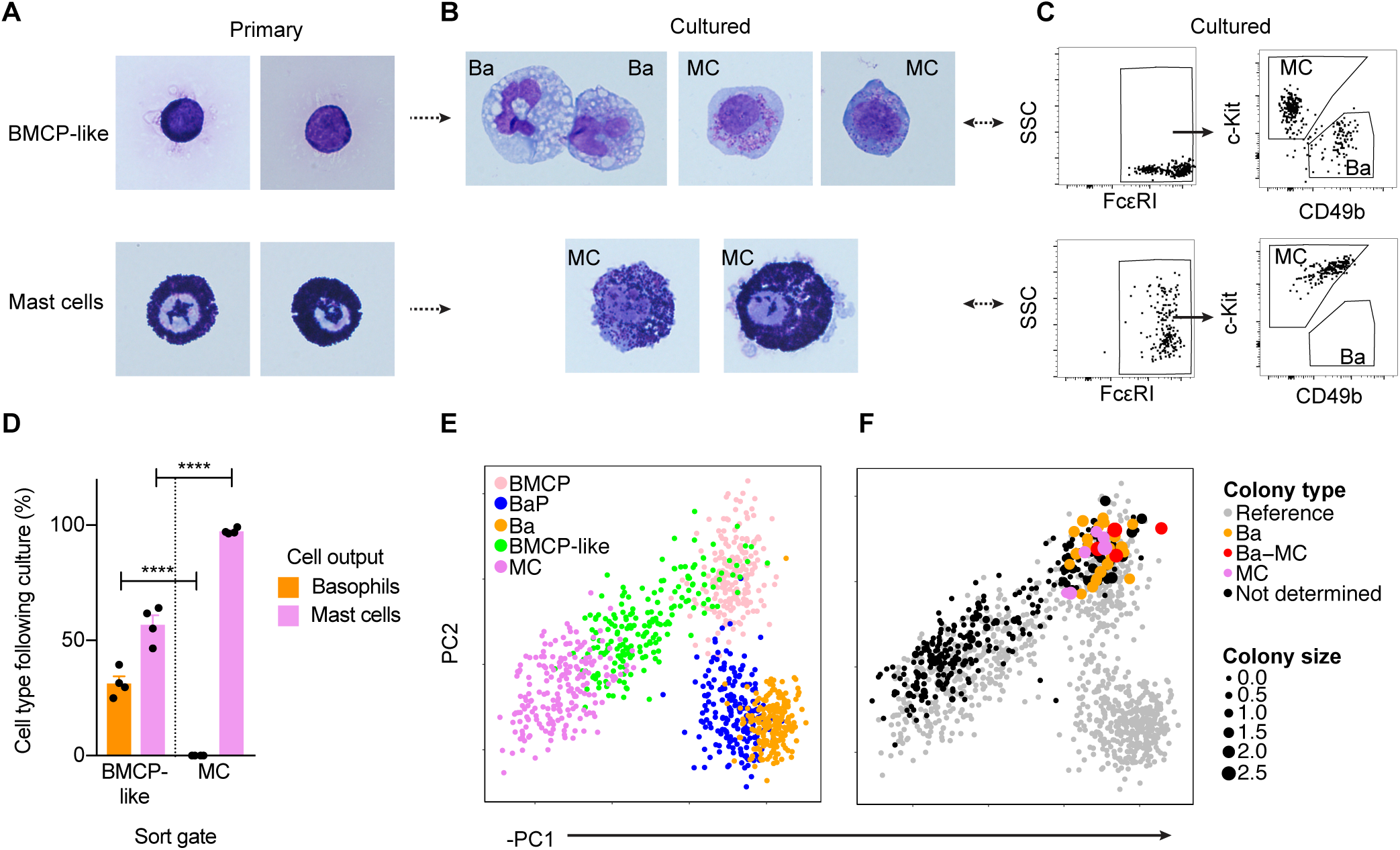
BMCP-like peritoneal cells exhibit potential to form basophils and mast cells. (A-B) May-Grünwald Giemsa staining of primary and in vitro cultured BMCP-like cells and mast cells extracted from the peritoneal cavity. Ba, basophil; MC, mast cell. Two or seven independent experiments revealed the morphology of primary BMCP-like cells and mast cells, respectively. (C) Flow cytometry gating strategy to identify basophils and mast cells cultured from primary BMCP-like cells and mast cells. (D) Quantification of cell type output following bulk-culture and flow cytometry analysis of BMCP-like cells and mast cells. Pooled data from 4 independent experiments per population are shown. The means and SEMs are shown. Unpaired two-tailed Student *t*-tests; \*\*\*\**P*<0.0001. (E) Principal component analysis of the flow cytometry reference dataset, provided in Figure 1C, colored by cell type. (F) Projection of index-sorted cells into the principal component space of the reference dataset. The point size represents log_10_-transformed colony size and the colors represent colony type following cell culture. Panel F shows data pooled from 2 independent experiments. The cells were cultured with IL-3 and stem cell factor.

We cultured the peritoneal cells to investigate whether the BMCP-like cell population exhibited capacity to generate basophils and mast cells. BMCP-like cells cultured with IL-3 and stem cell factor generated c-Kit^-^ FcεRI^+^ CD49b^+^ basophils and c-Kit^+^ FcεRI^+^ mast cells, whereas primary mast cells only displayed mast cell-forming capacity (Figure 4B-D).

By contrast to bulk cultured cells, only cell-fate assays performed at the single-cell level have the potential to reveal whether the BMCP-like population consists of bipotent progenitors. Therefore, single BMCP-like cells and mast cells were index sorted into individual wells, the resulting colony sizes were measured, and the colonies were subjected to flow cytometry analysis and cytochemical staining. To visualize the cell culture data, we first performed principal component analysis of the flow cytometry data presented in Figure 1C, henceforth referred to as the reference dataset (Figure 4E). We then projected the FACS index sort data onto the principal component space of the reference dataset, and plotted colony size and colony type data in the same embedding (Figure 4F). Analysis of colony sizes showed that colonies derived from BMCP-like cells were large, whereas cells along the mast cell trajectory exhibited reduced proliferation rate (Figure 4F). Notably, the cell-fate assays revealed that primary BMCP-like cells formed pure basophil colonies, pure mast cell colonies or mixed basophil-mast cell colonies (Figure 4F, S3A). Colonies derived from single mast cells were too small to analyze with flow cytometry. However, mast cells cultured in bulk remained mast cells as expected (Figure 4C-D, S3B-C).

We also cultured the BMCP-like peritoneal cells in erythroid-promoting conditions, as the early basophil-mast cell differentiation is closely linked to the erythrocyte trajectory.^2^ However, no erythroid output was observed (Figure S4), indicating that the BMCP-like cells indeed consisted of bipotent basophil-mast cell progenitors. Taken together, the cell culture assays revealed that the protein and gene expression analyses successfully predicted the differentiation state of the BMCP-like cell population. Our study therefore not only identifies a previously unknown bipotent peritoneal progenitor, but also provides comprehensive molecular profiles for this progenitor as well as the mast cell and basophil differentiation trajectories.

## Discussion

Here, we combine flow cytometry analysis, single-cell transcriptomics, and cell fate assays to chart the basophil and mast cell differentiation trajectories. Multicolor flow cytometry analysis reveals a developmental bifurcation with bipotent BMCPs and their progression into each respective mature lineage. High-coverage single-cell RNA sequencing allows us to generate a molecular map of the cell differentiation, and pseudotime ordering reveals dynamically regulated genes during development. We further demonstrate how flow cytometry and transcriptomics analysis can successfully predict cell-forming potential.

Single-cell transcriptomics coupled with index sorting of thousands of bone marrow HSPCs has previously been used to chart the erythrocyte and granulocyte-monocyte differentiation.^25, 26^ BMCPs represent a minor fraction of the bone marrow HSPCs, and capturing the early basophil-mast cell axis therefore requires analysis of tens of thousands of HSPCs.^3^ The early differentiation of progenitors with mast cell-forming capacity occurs in the bone marrow.^8^ However, full mast cell differentiation and maturation takes place at peripheral sites,^8^ and we therefore specifically sorted Lin^-^ c-Kit^+^ FcεRI^+^ cells extracted from the peritoneal cavity of mice to capture this process. Unlike cell extraction from tissues such as the intestine or skin, the isolation of cells from peritoneum does not require enzymatic digestion, thus minimizing external stimuli during cell processing. Basophil differentiation takes place in bone marrow, and we therefore analyzed basophils and their progenitors from this site. The combined peritoneal and bone marrow datasets provide a high-resolution map covering the BMCP bifurcation and the mast cell and basophil differentiation.

BMCPs have been described in the spleen and bone marrow,^3, 24^ and the presence of a bipotent progenitor population indicates that there is a close association between the basophil and mast cell differentiation trajectories. Recent data suggest that the erythroid axis is coupled with the basophil and/or mast cell fates.^2, 4, 27^ However, we did not observe erythrocyte-forming potential among 163 sorted BMCP-like cells in the peritoneum. In agreement with this, BMCPs in the spleen and bone marrow are unable to generate erythrocytes,^3, 24^ altogether suggesting that loss of erythrocyte-forming potential is a relatively early event along the differentiation trajectory from hematopoietic stem cells to basophil and mast cells. A similar differentiation process has been suggested in human hematopoiesis.^27^

Temporal ordering of the cells in the transcriptomic datasets allows exploration and verification of molecular processes in differentiating basophils and mast cells. We show that *Ndst2* (encoding *N*-deacetylase/*N*-sulphotransferase-2) is upregulated during differentiation from BMCPs to mature mast cells, and this was also associated with the appearance of numerous densely stained granules. In agreement with these findings, dense May-Grünwald Giemsa staining of the peritoneal mast cell granules requires sulphated heparin, which is dependent on *Ndst2* expression.^28^ Histamine is present in basophil and mast cells, and is quickly released upon cell activation. This potent mediator causes allergic reactions, and we show that the expression of the enzyme that catalyzes the histamine synthesis, histidine decarboxylase (*Hdc*), increased upon differentiation of both basophils and mast cells. Analysis of the single-cell transcriptomics data can also give insights into more complex regulatory processes. For example, autologous expression of integrin β7 is important for mast cell progenitor migration into tissues, such as lung and intestine.^29, 30^ Downregulation of integrin β7 on the cell surface is a hallmark of terminal mast cell differentiation,^31^ which we confirmed with flow cytometry analysis. However, we did not observe downregulation of the *Itgb7* gene expression during the transition from BMCPs to mast cells. Notably, integrins constitute αβ heterodimers when localized to the cell surface, and further investigation into the gene expression profile revealed decreased expression of *Itga4*, the binding partner of the integrin β7 subunit, upon differentiation. Thus, the loss of integrin α4 gene expression likely explains the loss of integrin β7 protein expression on the cell surface during the BMCP to mast cell transition.

During basophil differentiation, the transcription factors *Stat5b* and *Cebpa* are upregulated along the progression of pseudotime. The expression of C/EBPα is STAT5-dependent, and both genes are required for basophil formation.^24, 32^ Dynamic expression of transcription factors with currently unknown functions in basophil and mast cell differentiation was also recognized. For example, *Spi1*, which encodes PU.1, is upregulated during late basophil differentiation. It is known to be involved in neutrophil granulocyte maturation,^33, 34^ but the role of PU.1 in basophil differentiation is yet to be delineated. During mast cell differentiation, we describe the increase of the transcription factor *Meis2*. Primary mast cells from human skin express this transcription factor,^35^ but the potential function during mast cell differentiation is yet to be described. Thus, the datasets can be explored to identify previously unrecognized genes that may regulate basophil and mast cell differentiation.

Microarray analyses reported previously provide detailed gene expression patterns of mature hematopoietic cell populations, including bulk-sorted mature basophils and mast cells.^21^ We observed that differentiation into basophils and mast cells involves activation of mutually exclusive lineage programs. However, a small subset of the previously reported signature genes is not unique to mature cells, but can also be observed in bipotent progenitors. For example, we show that *Mcpt8* expression is not restricted to basophils but is also expressed by BMCPs. This in fact provides an explanation to a major conundrum in the field. Basophils, identified as *Mcpt8*-expressing cells, have been reported to exhibit potential to transdifferentiate into mast cells.^36^ Our results show that a more likely scenario is that a subset of the previously reported *Mcpt8*-expressing cells constitutes bipotent BMCPs that can give rise to mast cells.

Dimensionality reduction approaches are commonly applied to visualize single-cell transcriptomics data. Here, we show that the diffusion map embedding and PCA visualization successfully separate the basophil and mast cell differentiation trajectories based on the multidimensional flow cytometry data. We took advantage of the flow cytometry-based PCA visualization to interrogate single-cell fate assay data. The visualization showed that the in vitro proliferative capacity of the index-sorted peritoneal cells is quickly reduced as BMCP-like cells differentiate and enter the mast cell trajectory. In addition, only the most immature BMCP-like cells exhibit capacity to form basophil, mast cell, and mixed basophil-mast cell colonies. Taken together, dimensionality reduction techniques of flow cytometry data combined with cell fate assays provide new insights into basophil and mast cell differentiation.

In summary, here we have reported the generation of transcriptomic and flow cytometry data capturing the progression from bipotent progenitors toward basophil or mast cells. Our resource provides a detailed description of the expression changes occurring during this differentiation process at the single-cell level. A user-friendly interactive website has been created for the wider community to enable further exploration of the data.

## Authorship contributions

J.S.D. and W.W.Y.L. performed experiments; E.D. mapped sequencing data; F.K.H., J.S.D., and X.W. analyzed single-cell RNA sequencing data; J.S.D. analyzed flow cytometry and cell culture experiments; N.K.W. contributed to important discussions; I.K. created the web resource; B.G. and J.S.D. supervised the study; B.G. secured funding; F.K.H. and J.S.D drafted the manuscript; and all authors contributed to final version of the manuscript.

## Acknowledgements

We thank Chiara Cossetti, Gabriela Grondys-Kotarba, and Reiner Schulte at the Cambridge Institute for Medical Research Flow Cytometry Core for their assistance with cell sorting. J.S.D. is supported by funding from the Swedish Research Council, the Swedish Cancer Society, and Karolinska Institutet. Research in B.G.’s laboratory is supported by Bloodwise, Wellcome, CRUK, MRC and by core funding from Wellcome and MRC to the Wellcome-MRC Cambridge Stem Cell Institute. F.K.H. is funded by a MRC Physical Biology of Stem Cells PhD studentship and by part of a Wellcome Strategic Award (105031/D/14/Z) awarded to W. Reik, S. Teichmann, J. Nichols, B.D. Simons, T. Voet, S. Srinivas, L. Vallier, B.G. and J.C. Marioni.

## Supplementary Figure legends

**Figure S1.**
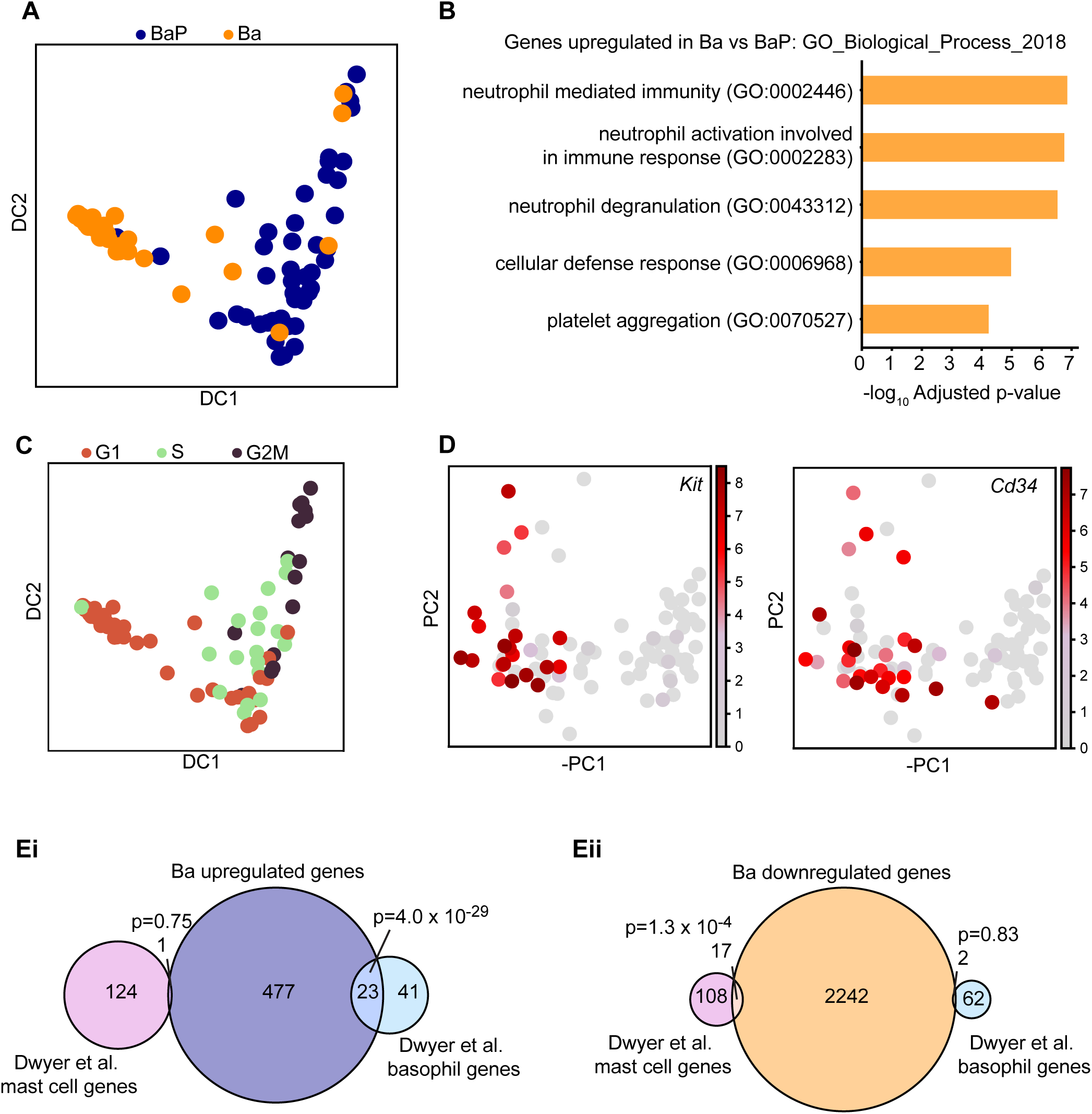
(A) Diffusion map of Ba and BaP cells colored by their phenotypic cell type. DC, diffusion component. (B) Top 5 GO Biological Process terms associated with the genes significantly upregulated in Ba cells compared to BaP cells, ranked by adjusted p-value. Benjamini-Hochberg correction for multiple hypotheses testing. (C) Diffusion map colored by computationally assigned cell cycle state. (D) Diffusion map colored by expression of specific genes. (E) Overlap of basophil (Ba) differentiation up- (i) or down- (ii) regulated genes with mast cell and basophil signature gene sets from Dwyer et al. Significance of overlap was tested using a hypergeometric test, with resulting p-values displayed in figure. The Venn diagrams show genes annotated in Ensembl genome build 81 and Dwyer et al.

**Figure S2.**
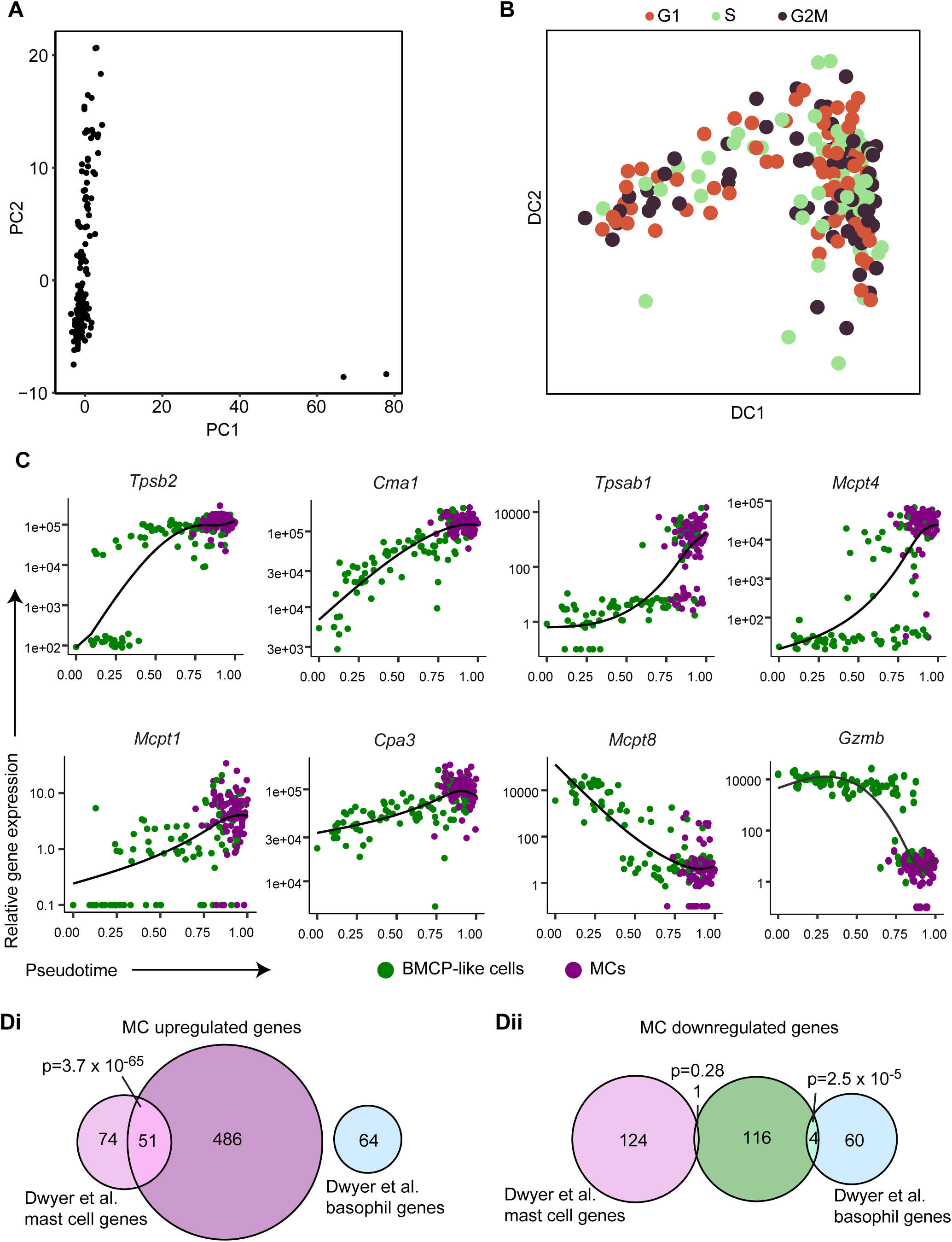
(A) PCA of peritoneal single-cell RNA-seq profiles showing outlier cells in PC1. PC, principal component. (B) Diffusion map dimensionality reduction colored by computationally assigned cell cycle state. DC, diffusion component. (C) Expression trends of specific genes along pseudotime. Splines were fitted using the monocle R package function. (D) Overlap of mast cell (MC) differentiation up- (i) or down- (ii) regulated genes with mast cell and basophil signature gene sets from Dwyer et al. Significance of overlap was tested using a hypergeometric test, with resulting p-values displayed in figure. The Venn diagrams show genes annotated in Ensembl genome build 81 and Dwyer et al.

**Figure S3.**
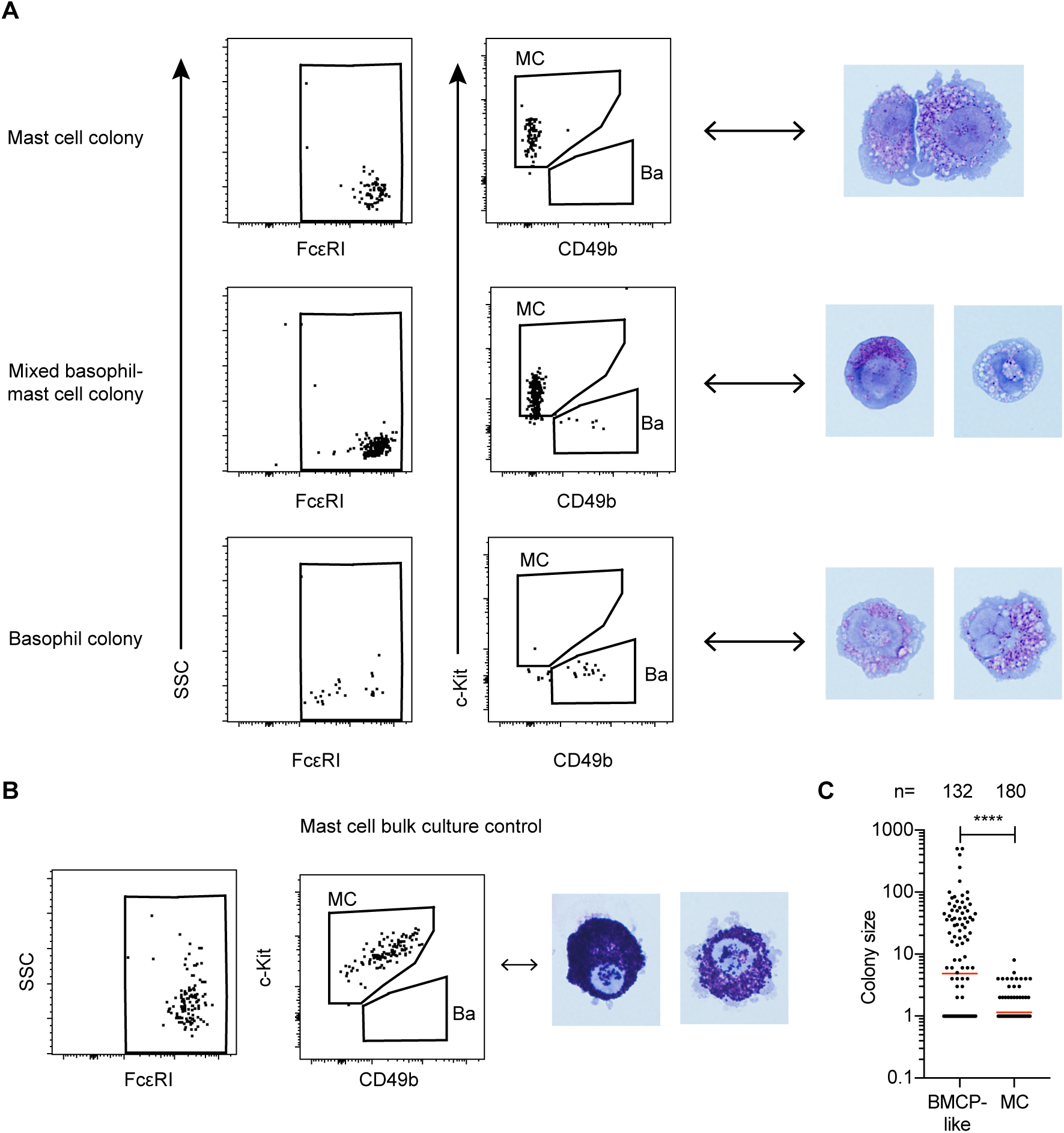
(A) Flow cytometry assessment of colonies derived from single BMCP-like peritoneal cells. The same colonies were stained with May-Grünwald Giemsa. (B) Cultured bulk-sorted peritoneal mast cells provided as reference to panel A. (C) Single-cells were sorted into individual wells and the colony size was determined after 7 days in culture. The numbers above each group represent the number of wells analyzed. Each dot represents one well. Wells in which no viable cells were found were scored as 1. The red lines represent geometric means. The data in panel C are pooled from 2 independent experiments. Two-tailed Mann-Whitney test; \*\*\*\**P*<0.0001. The cells were cultured with IL-3 and stem cell factor.

**Figure S4.**
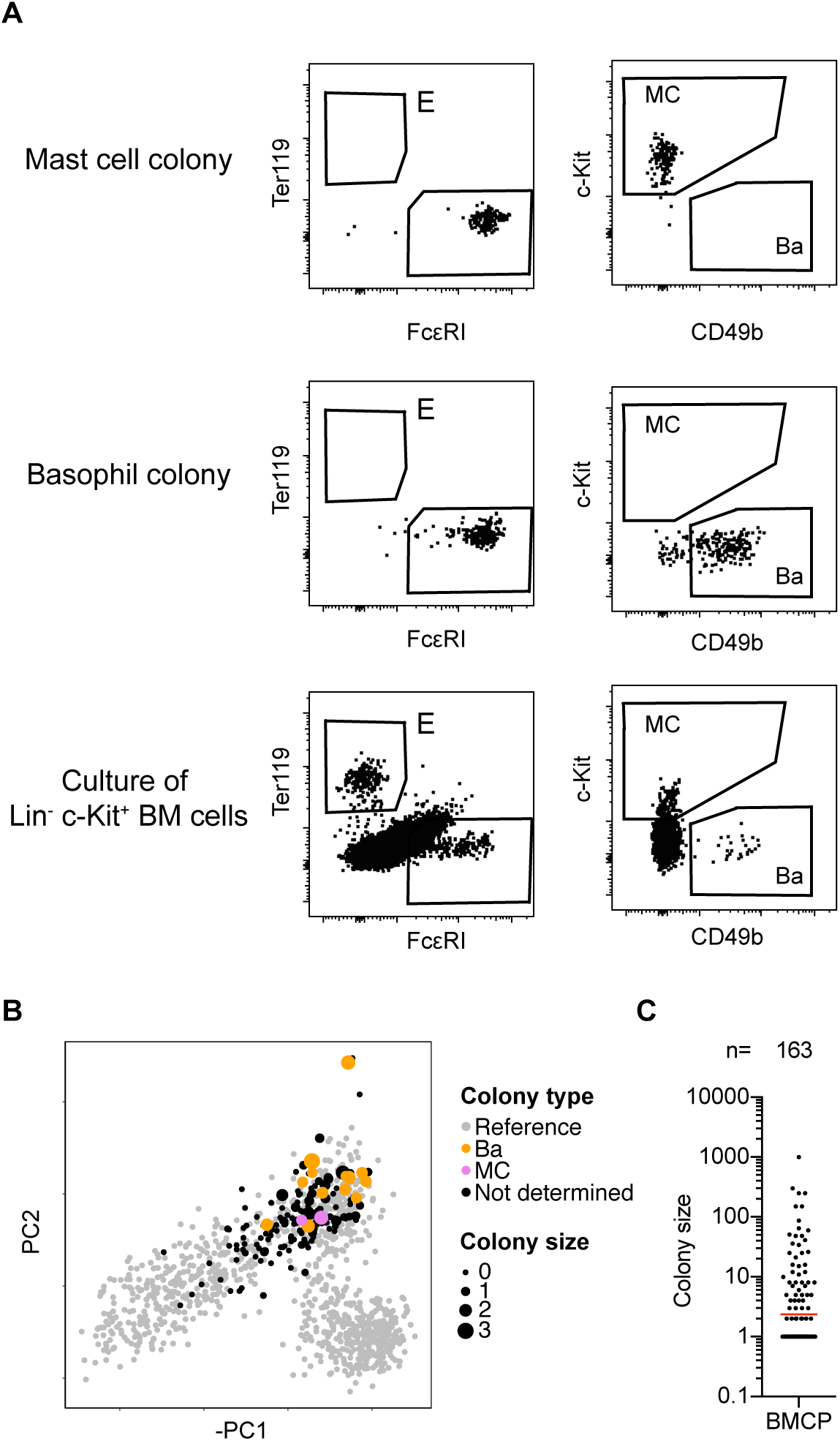
(A) Flow cytometry gating strategy for determining colony type following culture for 6 days in erythrocyte-promoting conditions. E, erythroid; MC, mast cell; Ba, basophil. Culture of bulk-sorted bone marrow (BM) Lin^-^ c-Kit^+^ cells served as positive control for erythroid-forming potential. (B) Colony type output of single index-sorted BMCP-like cells projected into the principal component space of the reference dataset. The point size represents log_10_-transformed colony size. (C) Single-cells were sorted into individual wells and the colony size was determined after 6 days in culture. The number represents the number of wells analyzed. Each dot represents one well. Wells in which no viable cells were found were scored as 1. The red line represents geometric mean. The data in panels B and C are pooled from 2 independent experiments.

